# Propranolol promotes bone formation and limits resorption through novel mechanisms during anabolic parathyroid hormone treatment in female C57BL/6J mice

**DOI:** 10.1101/2020.01.08.898320

**Authors:** Annika Treyball, Audrey C. Bergeron, Daniel J. Brooks, Audrie L. Langlais, Hina Hashmi, Kenichi Nagano, Deborah Barlow, Kathleen T. Nevola, Karen L. Houseknecht, Roland Baron, Mary L. Bouxsein, Anyonya R. Guntur, Katherine J. Motyl

## Abstract

Although the non-selective β-blocker, propranolol, improves bone density with PTH treatment in mice, the mechanism of this effect is unclear. To address this, we used a combination of *in vitro* and *in vivo* approaches to address how propranolol influences bone remodeling in the context of PTH treatment. In female C57BL/6J mice, intermittent PTH and propranolol had complementary effects in the trabecular bone of the distal femur and L5 vertebra, with combination treatment achieving micro-architectural parameters beyond that of PTH alone. Combined treatment improved the serum bone formation marker, P1NP, but did not impact other histomorphometric parameters relating to osteoblast function at the L5. *In vitro*, propranolol amplified the acute, PTH-induced, intracellular calcium signal in osteoblast-like cells. The most striking finding, however, was suppression of PTH-induced bone resorption. Despite this, PTH-induced receptor activator of nuclear factor kappa-B ligand (RANKL) mRNA and protein levels were unaltered by propranolol, which led us to hypothesize that propranolol could act directly on osteoclasts. Using *in situ* methods, we found *Adrb2* expression in osteoclasts *in vivo*, suggesting β-blockers may directly impact osteoclasts. Taken together, this work suggests a strong anti-osteoclastic effect of non-selective β-blockers *in vivo*, indicating that combining propranolol with PTH could be beneficial to patients with extremely low bone density.

## Introduction

The sympathetic nervous system (SNS) plays a critical role in the regulation of bone remodeling. The SNS suppresses bone formation and promotes receptor activator of nuclear factor kappa-B ligand (RANKL)-mediated osteoclast recruitment, resulting in low trabecular bone volume fraction in mice ^(1,2)^. Consistent with this, tyrosine hydroxylase, one of the enzymes involved in norepinephrine production, is present in nerves within bone, marrow, and periosteum ^(3)^. The major adrenergic receptor mediating the downstream effects of norepinephrine appears to be the β2AR in mice ^(1)^. Deletion of β2AR, or treatment with the non-selective βAR antagonist (β-blocker) propranolol, prevents bone loss in situations of stress, antipsychotic and antidepressant treatment, and in other situations in which sympathetic signaling to bone may be elevated ^(1,4–8)^. The majority of studies point toward a mechanism of β2AR suppressing bone formation directly in the osteoblast, and activating bone resorption indirectly, through the RANKL/OPG pathway ^(1,2,9)^. Indeed, mice with an osteoblast-specific deletion of *Adrb2* have a high bone density phenotype as adults, with increased bone formation and reduced *Rankl* expression ^(10)^. However, some studies have found evidence for direct effects of β2AR signaling in osteoclasts ^(11,12)^, suggesting the downstream effects of elevated SNS activity may be more complex than generally accepted.

Despite this complexity, identifying osteoporosis treatment strategies that modulate sympathetic signaling remains clinically useful. β-blockers are one such group of drugs that have been shown to reduce fracture risk and increase BMD in patients ^(13)(14)^. Furthermore, rodent studies have shown that combining β-blockers with teriparatide (intermittent truncated parathyroid hormone, PTH) may further promote bone density. Evidence from ovariectomized mice suggests that combining PTH treatment with propranolol increases bone mineral density beyond the levels achieved by PTH alone, and histomorphometric analyses indicated that improvement was largely due to increased bone formation and osteoblast number ^(15)^.

On a cellular level, evidence from *in vitro* and *in vivo* studies suggest that PTH efficacy is dependent upon the presence of the β2AR. Deletion of *Adrb2*, the gene that encodes β2AR, prevents the anabolic effect of intermittent PTH in young and aged mice, suggesting some β2AR signaling is required ^(16)^. One mechanism of this may be through β2AR signaling allowing G-protein βγ (Gβγ) subunit to bind endosomal PTH1R, which sustains cAMP levels leading to enhanced mineralization in osteoblasts ^(17)^. cAMP signaling, however, is not the only avenue for PTH effects to be transduced to the cell ^(18)(19)^. PTH1R also signals through phospholipase C to increase Ca^2+^ release from intracellular stores, as well as through arrestin-mediated mechanisms ^(18)(19)^. There is an established role for intracellular Ca^2+^ signaling in the regulation of osteoblast differentiation through the calmodulin/CamKII pathway regulating AP-1 and CREB/ATF4 transcription factors ^(20)^. It has been shown that silencing of *Adrb2* can increase pCREB ^(21)^, but it remains unknown whether the combination of PTH treatment with β2AR antagonists would impact these pathways.

To further investigate the cellular and systemic effects of co-modulation of PTH1R and β2AR, we performed a series of *in vitro* and *in vivo* assays pharmacologically targeting these mreceptors. A pharmacological approach was chosen so that any advantages of combination treatments may eventually be translated to humans. Briefly, we determined that the combination of PTH and propranolol *in vivo* increased bone volume fraction in the distal femur in part by increasing markers of bone formation, which may be related to the β-blocker propranolol enhancing PTH-induced Ca^2+^ signaling in osteoblast-like cells *in vitro*. More striking, however, was that propranolol prevented PTH-induced bone resorption, but did not impact PTH-induced changes in RANKL or OPG pathway members. Rather, we identified that this effect may be through direct anti-osteoclastogenic effects of propranolol since *Adrb2* is expressed in osteoclasts *in vivo*. In all, these findings suggest that modifications to PTH therapy that mimic outcomes from propranolol treatment, or simply combining PTH with β-blockers, may be a useful approach to minimizing resorption and promoting net bone accrual.

## Results

### Propranolol improved total BMD in the presence of PTH

To test whether propranolol would improve PTH-induced bone formation, we treated mice with either vehicle, PTH, propranolol, or PTH + propranolol from 16-20 weeks of age. Neither body weight, fat mass, nor fat-free mass was altered in any of the treatment groups, suggesting that any bone changes would be independent of differences in loading (Table I). As expected, PTH had a significant main effect on total aBMD and aBMC, as well as on femoral aBMD (Table I). Although propranolol only had a significant main effect on total aBMD, all bone parameters measured by DXA were highest in the PTH + propranolol group (Table I). Interestingly, co-treatment with PTH and propranolol increased total aBMD beyond that caused by PTH or propranolol alone.

**Table I.**
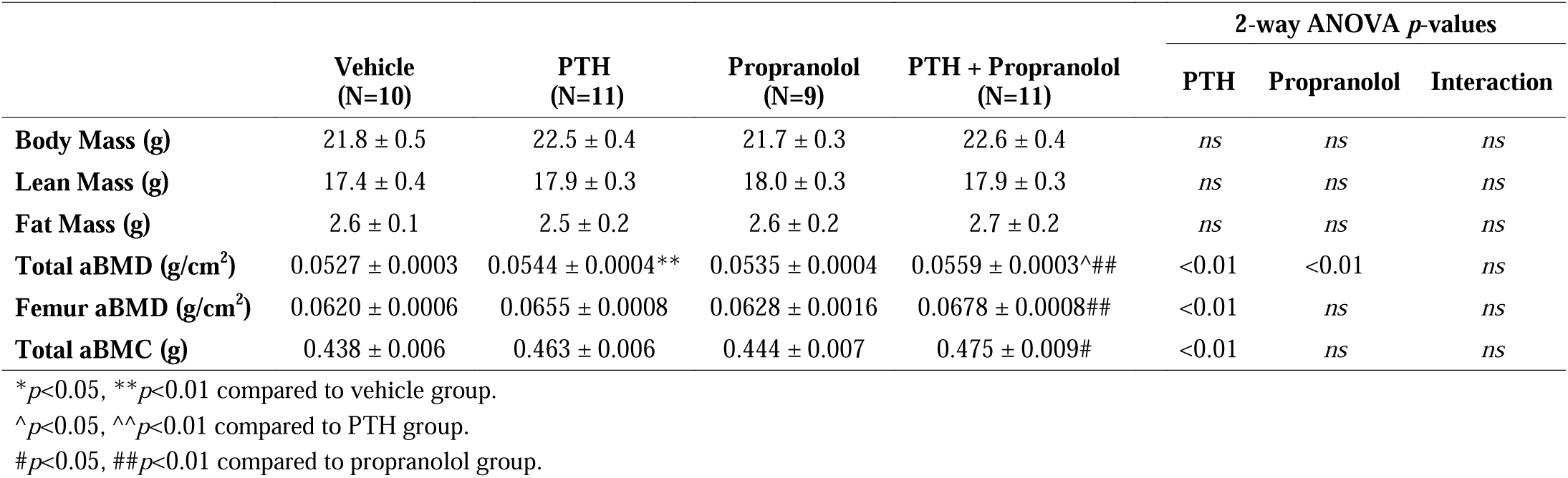
Body composition and areal bone parameters from mice treated with PTH and/or propranolol.

### PTH and propranolol had site-specific effects on trabecular bone microarchitecture

To investigate the impact of PTH and propranolol treatment on bone microarchitecture, we performed µCT on L5 vertebrae and femurs. Propranolol and PTH both independently improved trabecular BV/TV of the L5 vertebra, but the combination of PTH and propranolol improved BV/TV and BMD above and beyond that of either treatment alone (Figure 1). This is likely due to a combined effect of increased trabecular thickness and increased trabecular number, the latter of which was only significantly increased by PTH when propranolol was present. Consistent with this, PTH and propranolol together significantly reduced BS/BV ratio compared to propranolol alone (Figure 1).

**Figure 1.**
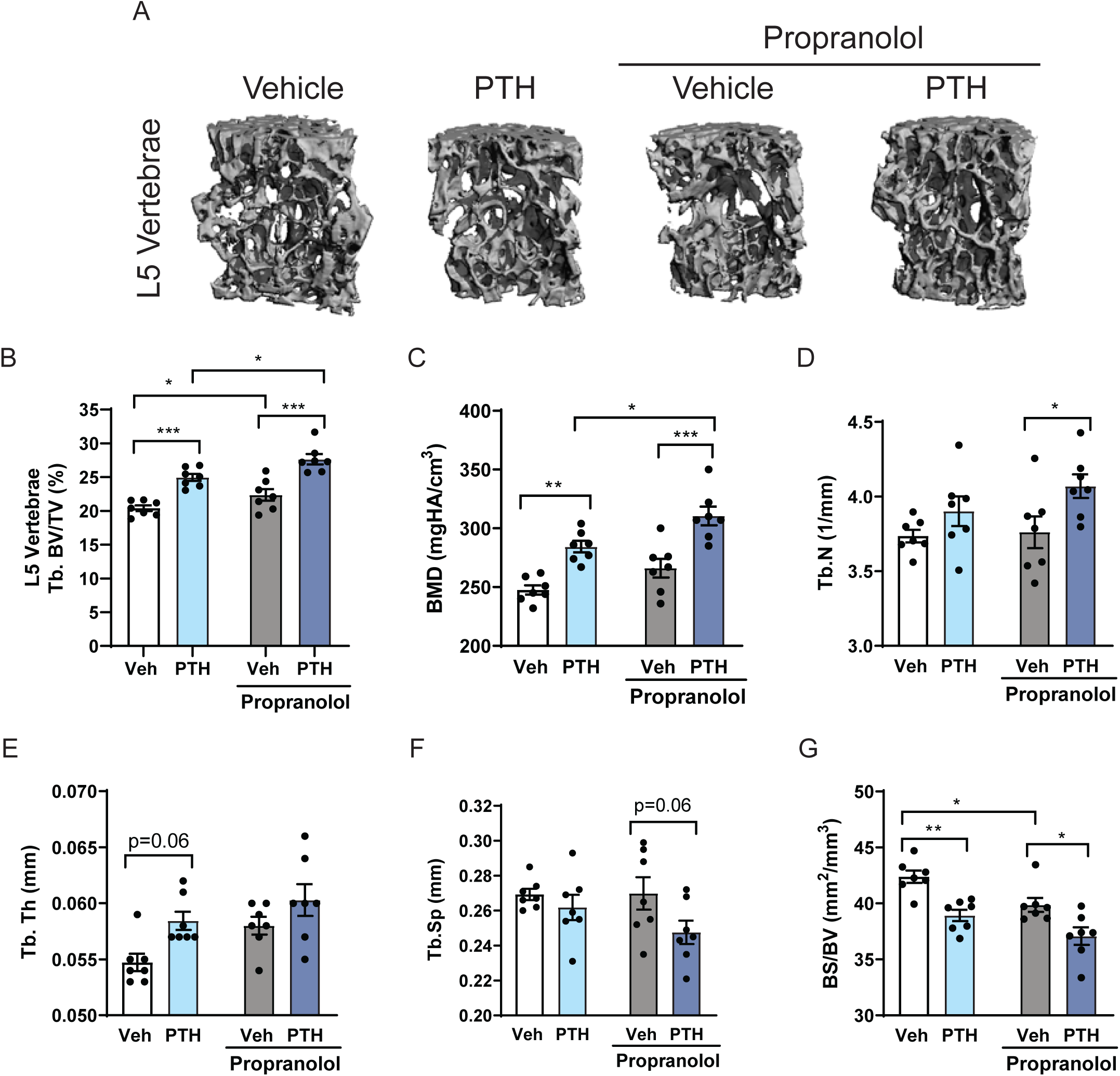
Propranolol improved the positive effects of PTH on trabecular bone in the L5 vertebrae. Mice were treated with vehicle (white), 80 µg/kg PTH (light blue), 0.5 mg/ml propranolol (gray), or PTH and propranolol (dark blue) for 4 weeks, from 16-20 weeks of age. (A) Representative µCT images from L5 vertebrae. (B-G) Trabecular bone volume fraction (Tb. BV/TV), bone mineral density (BMD), number (Tb.N), thickness (Tb.Th), separation (Tb.Sp), and bone surface/bone volume (BS/BV). Bars represent mean ± standard error. \**p*<0.05, \*\**p*<0.01 by Holm-Sidak *post hoc* test after a significant two-way ANOVA.

In the distal femur, we focused our examination of PTH and propranolol effects on two distinct sites, the primary and secondary spongiosa (Figure 2). In the secondary spongiosa, PTH increased Tb. BV/TV, BMD, and Conn.D and these parameters were all increased further by combined treatment with propranolol (Figure 2B-D). Although some parameters (Tb.N, Tb.Th, Tb.Sp, and BS/BV) were not altered by PTH alone, the combination reduced SMI, Tb.Sp, and BS/BV, while significantly increasing Tb.N and Tb.Th (Figure 2E-I). The primary spongiosa also had striking changes from PTH, including increased BV/TV, BV and BMD (Figure 2J-L), but these were not exacerbated or diminished by combination PTH and propranolol treatment. Tissue mineral density (TMD), a measurement of the density of the bone itself (not including any marrow), was suppressed with PTH treatment, suggesting that mineral deposition may be compromised (Figure 2M). Interestingly, however, propranolol significantly elevated TMD in PTH-treated mice such that it was not different from vehicle-treated (Figure 2M). This indicates the quality of mineralization in the primary spongiosa during combination PTH and propranolol treatment may be improved compared to PTH alone.

**Figure 2.**
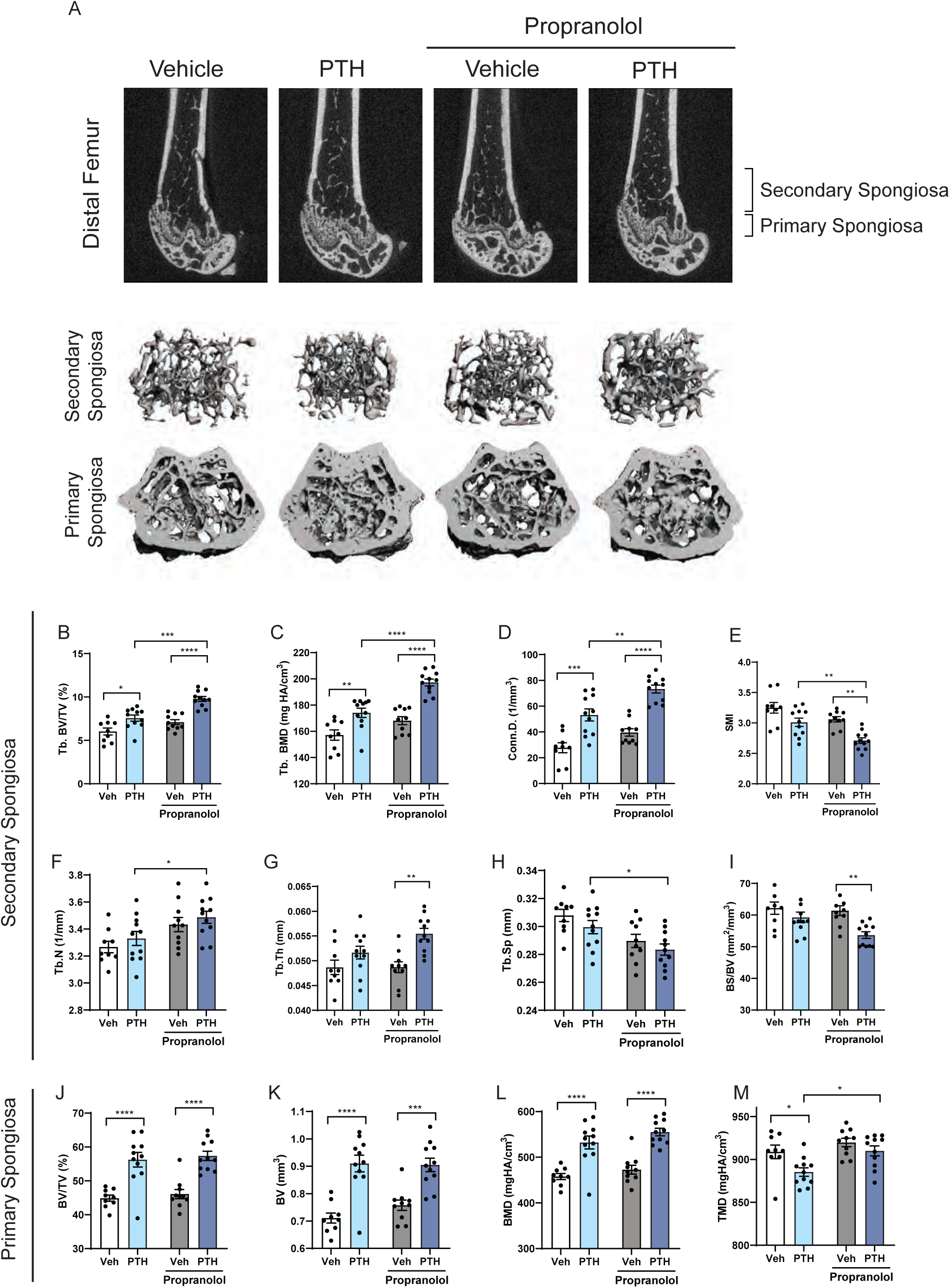
Trabecular bone in the femur secondary spongiosa is enhanced by combination PTH and propranolol treatment. Mice were treated with vehicle (white), 80 µg/kg PTH (light blue), 0.5 mg/ml propranolol (gray), or PTH and propranolol (dark blue) for 4 weeks, from 16-20 weeks of age. (A) Representative 2D (top) and 3D (bottom) images of trabecular micro architecture in the primary and secondary spongiosa of the distal femur. (B-I) Trabecular microarchitectural parameters from the secondary spongiosa. (J-K) Volumetric and densitometric measurements from the primary spongiosa. Bars represent mean ± standard error. \**p*<0.05, \*\**p*<0.01, \*\*\**p*<0.001, \*\*\*\**p*<0.0001 by Holm-Sidak *post hoc* test after a significant two-way ANOVA.

### Cortical bone microarchitecture

PTH significantly increased cortical area (Ct.Ar) and polar moment of inertia (pMOI) in the femur midshaft, but did not significantly impact marrow area (Ma.Ar), total area (Tt.Ar), cortical thickness (Ct.Th), tissue mineral density (TMD) or porosity (Figure 3 A-H). Although propranolol treatment did not impact these parameters on its own, propranolol significantly improved the effect of PTH on the cortical bone such that Ct.Ar, Ct.Ar/Tt.Ar, Ct.Th and TMD were elevated in the PTH + propranolol group compared to propranolol alone (Figure 3A, D, E, and F). The increased Ct.Ar/Tt.Ar with combination treatment is most likely due to a reduction in Ma.Ar, because Tt.Ar was clearly unchanged (Figure 3B-D).

**Figure 3.**
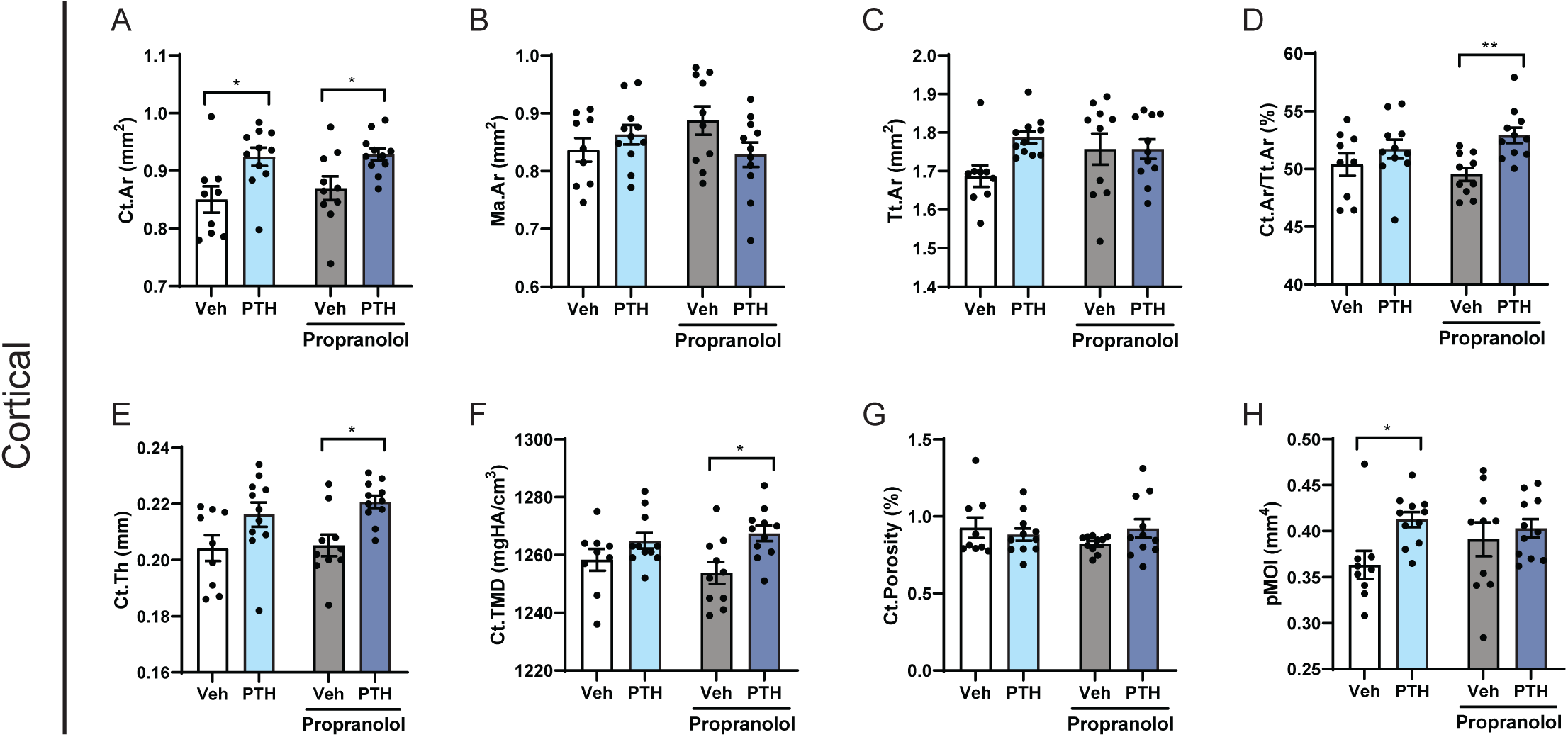
Propranolol improved cortical thickness and density in PTH-treated mice. Mice were treated with vehicle (white), 80 µg/kg PTH (light blue), 0.5 mg/ml propranolol (gray), or PTH and propranolol (dark blue) for 4 weeks, from 16-20 weeks of age. (A-H) Cortical bone microarchitectural parameters, cortical area (Ct.Ar), marrow area (Ma.Ar), total area (Tt.Ar) cortical area fraction (Ct.Ar/Tt.Ar), cortical thickness (Ct.Th), cortical tissue mineral density (Ct.TMD), cortical porosity, and polar moment of inertia (pMOI) were analyzed at the midshaft of the femur. Bars represent mean ± standard error. \**p*<0.05, \*\**p*<0.01 by Holm-Sidak *post hoc* test after a significant two-way ANOVA.

### Propranolol promoted bone formation in PTH-treated mice

To determine whether co-treatment with propranolol modulated the effect of PTH on bone formation, we performed serum, histomorphometric, and mRNA analyses to evaluate bone remodeling activity. As expected, PTH increased the serum marker of bone formation, P1NP, and this was further increased by co-treatment with PTH and propranolol (Figure 4A). Histomorphometric analyses in the L5 vertebrae indicated a significant effect of PTH by 2-way ANOVA on MAR and BFR parameters (Figure 4B, C). Similarly, osteoblast parameters were elevated by PTH, but were not significantly modulated by the combination with propranolol. Propranolol did not have a significant main effect on these parameters (except to decrease the N.Ob/B.Pm) and, when combined with PTH, did not increase these indices beyond the level of PTH alone (Figure 4B, C). Propranolol did, however, reduce the amount of OS/BS in PTH-treated mice (Figure 4C).

**Figure 4.**
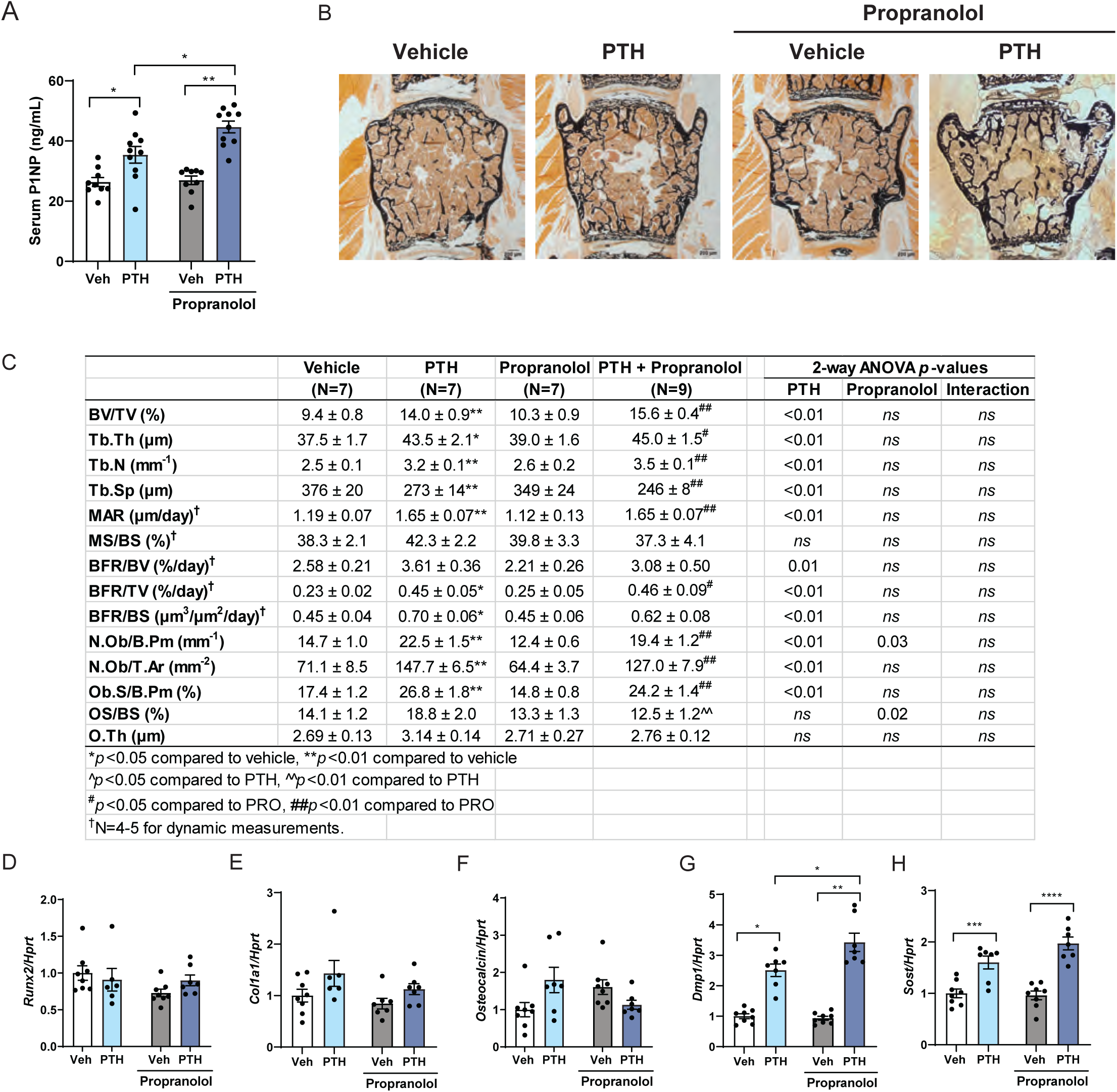
Propranolol improved bone formation markers in PTH-treated mice. Mice were treated with vehicle (white), 80 µg/kg PTH (light blue), 0.5 mg/ml propranolol (gray), or PTH and propranolol (dark blue) for 4 weeks, from 16-20 weeks of age. (A) Serum P1NP was measured by ELISA. (B-C) Static and dynamic histomorphometry representative images and quantification of architectural, bone formation and osteoblast parameters measured in L5 vertebrae. (D-H) Gene expression of markers of bone formation and osteoblast maturity were analyzed in the tibia and normalized by the non-modulated housekeeping gene, *Hprt*. Bars represent mean ± standard error. \**p*<0.05, \*\**p*<0.01 by Holm-Sidak *post hoc* test after a significant two-way ANOVA.

In general, these findings contrast the serum P1NP finding, suggesting other sites may have elevated bone formation with the combination treatment. Therefore, we examined mRNA markers of osteoblastogenesis and osteocyte maturation in a separate site, the tibia. Although we did not see significant differences in *Runx2, Col1a1*, or *Osteocalcin* expression (Figure 4D-F), we did observe differences in late osteoblast and osteocyte markers. First, dentin matrix acidic phosphoprotein 1 (*Dmp1*) expression was increased with PTH treatment, but there was also a significant interaction (p=0.01) such that PTH-induced *Dmp1* expression was higher in the propranolol treated group, while propranolol alone had no effect in the absence of PTH (Figure 4G). Surprisingly, intermittent treatment with PTH induced *Sost* expression, in contrast to previously reported effects with chronic elevation, and *Sost* was elevated to a greater degree in propranolol treated mice (Figure 4H) ^(22)^. Although serum P1NP is elevated, changes in bone formation (histological and mRNA-based) do not convincingly explain the improved BV/TV in mice treated with PTH and propranolol.

Recent studies have suggested PTH may suppress bone marrow adipose tissue (BMAT) as a means or consequence of promoting bone formation. Although marrow adipocyte size was not influenced by PTH, there was a subtle but significant interaction effect (p<0.05) of PTH and propranolol in both adipocyte volume / total volume (AV/TV) and adipocyte number parameters. In particular, PTH tended to reduce marrow adipocyte numbers in mice not treated with propranolol (p=0.08) but this effect was blunted in propranolol-treated mice (Figure S1).

### PTH-induced intracellular calcium signaling is enhanced by propranolol

Next, we sought to test the impact of propranolol and PTH directly in osteoblasts by measuring the concentration-dependence of the intracellular Ca^2+^ response to PTH in MC3T3-E1 cells differentiated for 7 days (Figure 5). PTH induced a measurable Ca^2+^ signal at concentrations as low as 10 nM and the peak response was observed at 1 µM. Using a log transformation, we calculated the concentration with 50% of maximal excitation to be 66.6 nM (Figure 5A), consistent with previously published reports (published iCa^2+^ EC_50_ = 50 nM with dynamic range 5-500 nM)^(23)^. Therefore, we used a concentration of 100 nM in our studies, to be within the dynamic range of the concentration-response curve, but also within the concentrations used in previous publications demonstrating functional effects of PTH on osteoblasts^(24)^. Next, we found that pre-treatment with propranolol significantly increased the intracellular Ca^2+^ signal induced by PTH (Figure 5B, C). This is in contrast to treatment with propranolol alone, which did not induce any intracellular Ca^2+^ signal (Figure 5B).

**Figure 5.**
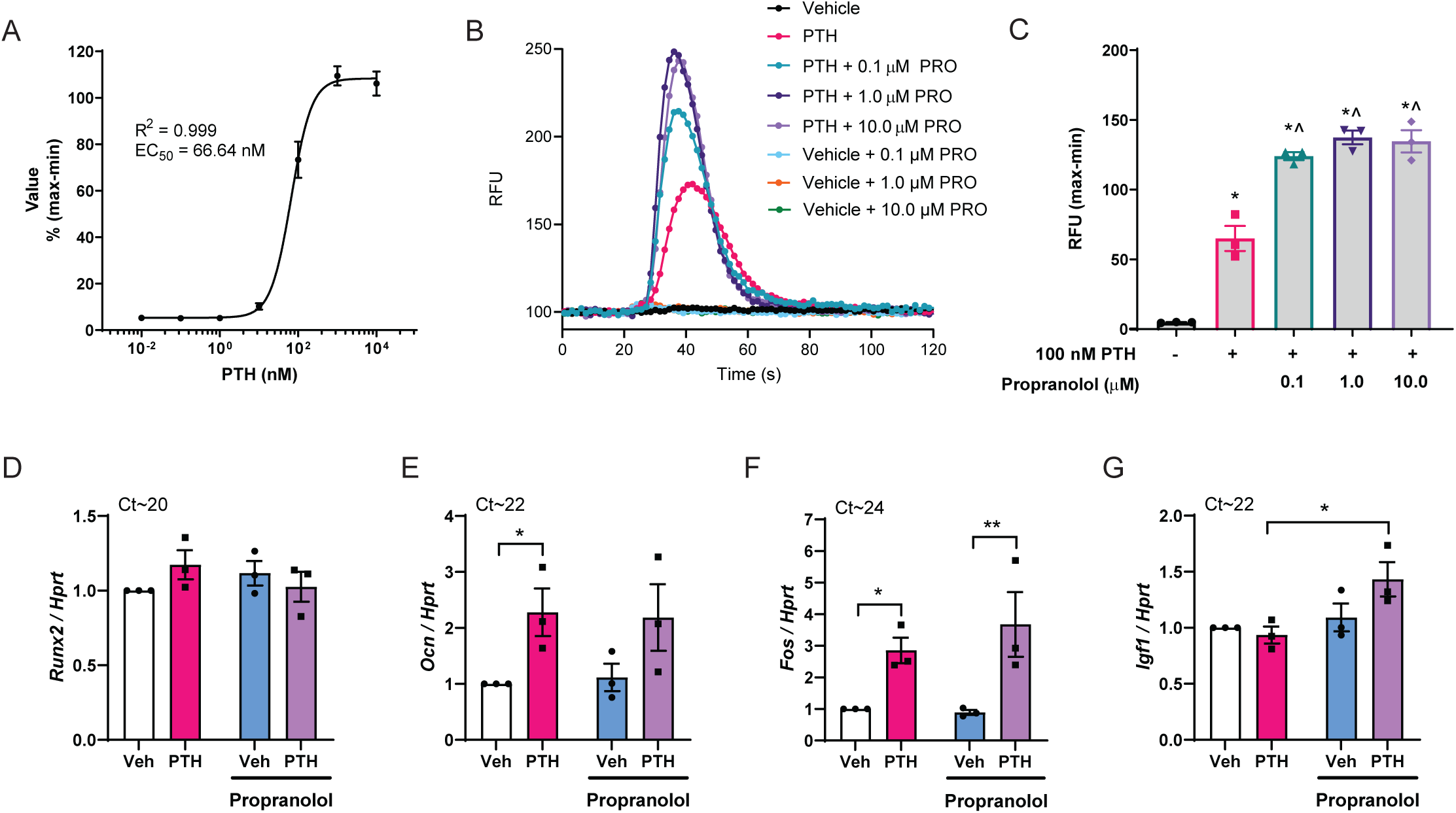
β-blocker propranolol potentiated osteoblast intracellular Ca^2+^ and *Igf1* expression. MC3T3-E1 cells were differentiated to day 7 in osteogenic media. (A) Fluorescence ratio indicative of intracellular Ca^2+^ concentrations was expressed as a percentage of the maximum level. (B) Representative traces of 100 nM PTH or vehicle-induced fluorescence over time with pretreatment of vehicle, or 0.1, 1 or 10 µM propranolol. Note: vehicle + propranolol pretreatment did not induce a significant signal, and traces are only slightly visible behind vehicle-only trace. (C) Quantification of the maximum-minimum PTH-induced Ca^2+^ fluorescence after pretreatment with vehicle or propranolol. One representative of three experiments (each performed in triplicate) is shown. \**p*<0.05 compared to vehicle-treated (black), ^*p*<0.05 compared to 100 nM PTH-treated (pink) by Tukey’s *post-hoc* test after significant one-way ANOVA. (D-K) Gene expression of *Runx2, Ocn, Fos*, and *Igf1* in MC3T3-E1 cells at day 7 after 1 hour treatment with vehicle (white), 100 nM PTH (pink), 1 µM propranolol (blue) or 100 nM PTH + 1 µM propranolol (purple). Genes are normalized to non-modulated housekeeping gene, *Hprt*. Each data point represents the mean from an independent experiment, performed in triplicate. Expression levels in vehicle-treated cells from each experiment were set to 1. Approximate Ct values of vehicle-treated groups are shown above each graph. Bars represent mean of the three independent experiments ± SEM. \**p*<0.05, \*\**p*<0.01 by Holm-Sidak *post hoc* test after a significant two-way ANOVA.

Long-term PTH treatment does not promote mineralization *in vitro*, so to get a sense whether combined PTH and propranolol treatment had any functional consequence in osteoblasts, we examined gene expression of relevant receptors (Figure S2) and osteoblast markers after acute treatment with PTH, propranolol or both (Figure 5D-G). Although there were no differences in early osteoblastogenic markers *Runx2* (Figure 5D) and *Sp7* (not shown), PTH induced *Osteocalcin* expression (main effect of PTH *p*=0.016) similarly in both vehicle and propranolol treated cells but only the group treated with PTH alone (no propranolol present) reached pairwise statistical significance (Figure 5E). Because intracellular Ca^2+^ has been shown to modulate AP-1 family transcription factors, we examined gene expression of *Fos*. Interestingly, the PTH-induced increase in *Fos* was higher when propranolol was present (Figure 5F), consistent with propranolol promoting PTH-induced intracellular Ca^2+^ levels. One established mechanism of osteoanabolism from PTH is through increased expression and autocrine signaling of IGF-1^(24)^. We found a significant increase in *Igf1* expression only in the PTH and propranolol co-treated group (Figure 5G). These findings suggest propranolol may improve bone formation from PTH in part through increasing intracellular Ca^2+^ signaling and promoting expression of *Igf1*.

### PTH-induced osteoclast activity was blocked by propranolol

Since PTH is known to enhance resorption, we next investigated whether propranolol mitigates resorption measurements *in vivo*. PTH increased expression of *Ctsk* and *Trap5*, as well as the serum protein levels of the resorption marker CTx (Figure 6A-C). Interestingly, propranolol prevented the PTH-induced increases in *Trap5* expression and serum CTx levels, suggesting propranolol prevents PTH-induced resorption (Figure 6B, C). Consistent with the literature, histomorphometry indicated that PTH increased the total number of osteoclasts (N.Oc/T.Ar, Figure 6D). Propranolol alone did not have any impact on the number of osteoclasts or the other osteoclast parameters (Figure 6D). When combined, however, the number of osteoclasts on the bone surface and within the total area measured (N.Oc/B.Pm and N.Oc/T.Ar, respectively) and the osteoclast surface fraction (Oc.S/BS) was lower in mice treated with PTH and propranolol compared to those treated with PTH alone (Figure 6D), interaction p≤0.02). Notably, the PTH-induced increase in N.Oc/T.Ar was completely blocked by co-treatment with propranolol (*p*<0.01, Figure 6D). All three lines of evidence related to osteoclasts (serum, mRNA, and histomorphometry) point toward a strong effect of propranolol to reduce PTH-induced bone resorption.

**Figure 6.**
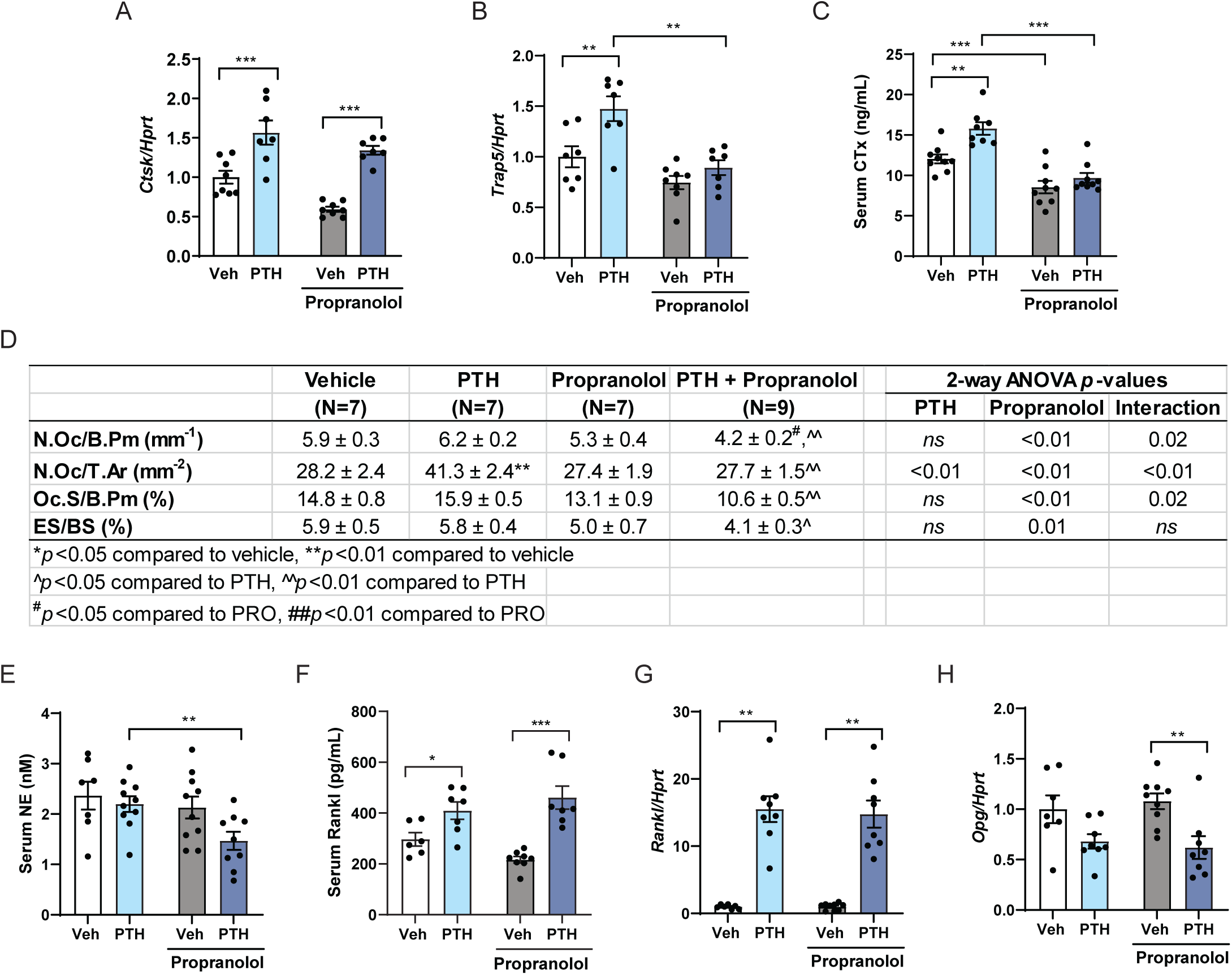
Propranolol prevented PTH-induced resorption but not RANKL induction. Mice were treated for 4 weeks (A-F) with vehicle (white), 80 µg/kg PTH (light blue), 0.5 mg/ml propranolol (gray), or PTH and propranolol (dark blue). (A-B) Gene expression of *Ctsk* and *Trap5* was analyzed in the whole tibia and normalized by the non-modulated housekeeping gene, *Hprt*. (C) Serum CTx was measured by ELISA. (D) Histomorphometric analyses of osteoclast and resorption parameters were measured in L5 vertebrae. (E) Serum norepinephrine (NE) was measured by LC/MS/MS. (F) Serum Rankl was measured by ELISA. (G-H) Mice were treated with PTH and/or propranolol for 5 days and cortical shell was collected 1 hour after the last PTH or vehicle dose. Gene expression levels of *Rankl* and *Opg* were normalized to *Hprt*. Bars represent mean ± standard error. \**p*<0.05, \*\**p*<0.01, \*\*\**p*<0.001 by Holm-Sidak *post hoc* test after a significant two-way ANOVA.

Propranolol can centrally suppress sympathetic activity, therefore we measured norepinephrine (NE) levels in serum as well. While neither PTH nor propranolol independently modulated NE, the combination treatment resulted in significantly lower circulating NE compared to PTH alone (Figure 6E), suggesting a systemic lowering of sympathetic activity by propranolol might be responsible for the suppressed resorption. Because the current understanding of the mechanism of NE signaling-induced osteoclastogenesis is through the modulation of *receptor activator of nuclear factor kappa-B ligand* (*Rankl*) expression in the osteoblast, we measured circulating RANKL and found that, to our surprise, propranolol had no effect on PTH-induced RANKL levels (Figure 6F). Furthermore, although PTH induced *Rankl* gene expression in cortical bone as expected, propranolol did not have any blunting effect on *Rankl* (Figure 6G), but did lower *Opg* (Figure 6H), suggesting other mechanisms were responsible for the reduced osteoclast numbers and CTx levels in PTH and propranolol treated mice.

Previous *in vitro* work has suggested βARs may play a direct role in osteoclasts ^(11,12)^, but this has not been confirmed *in vivo*. To test whether β2AR may be directly responsible for the anti-osteoclastic effects of propranolol, we first tested marrow and serum concentrations of propranolol achieved after dosing mice in the same manner. Propranolol concentrations achieved in the serum of mice dosed in drinking water were 14.6 nM, while bone marrow concentrations were 107.7 nM (Figure 7A). Within each mouse, approximately ten times more propranolol was present in marrow compared to serum, suggesting osteoclasts are likely to have significant propranolol exposure. We next examined whether *Adrb2* expression was present in osteoclasts *in vivo* using *in situ* hybridization. First, we used probes specific to *Acp5*, the gene encoding TRAP5, to identify multinucleated osteoclasts (Figure 7B). Next, we co-labeled with probes specific to *Adrb2* and found levels of *Adrb2* expression in osteoclasts comparable to that of other cells in the marrow (Figure 7C). Collectively, these findings suggest that lowered NE levels combined with direct suppression of osteoclasts by propranolol may explain the reduced resorption and improved micro-architectural parameters in mice co-treated with PTH and propranolol.

**Figure 7.**
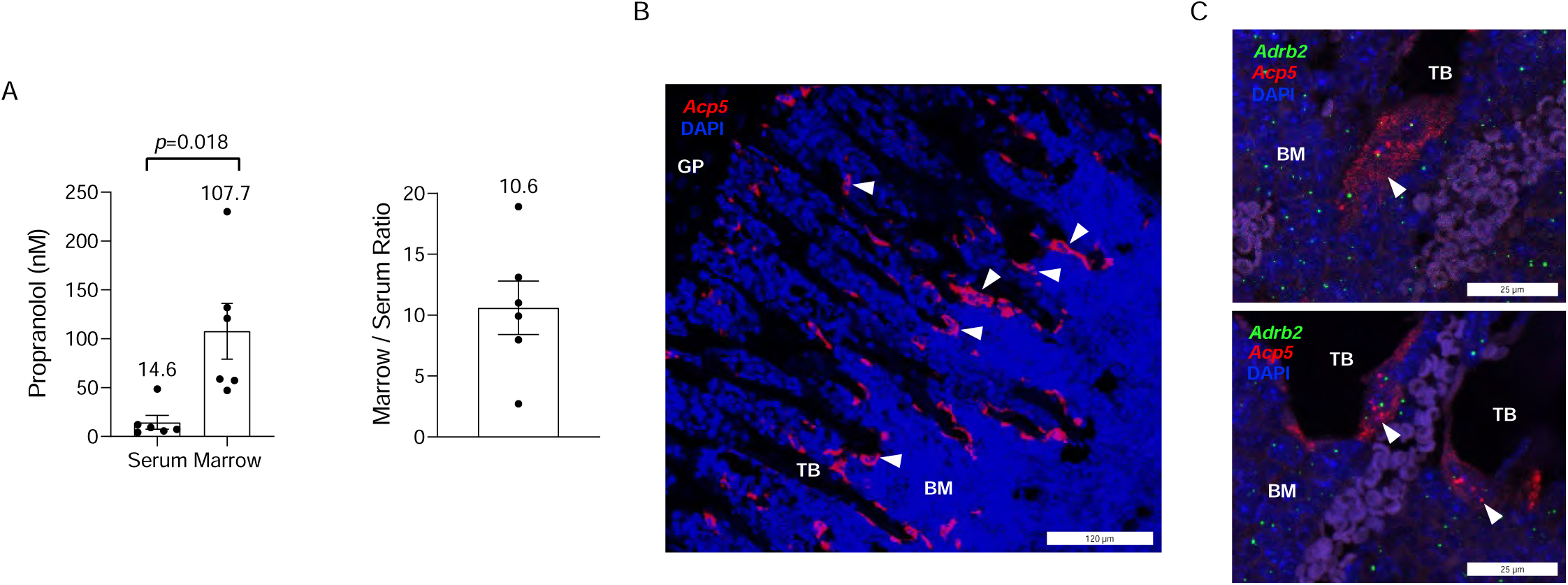
Propranolol is present in marrow, where *Adrb2* is expressed in osteoclasts *in vivo*. (A) *Left:* Serum and marrow concentrations of propranolol after four days of treatment with 0.5 mg/mL in the drinking water, measured by LC/MS/MS. *p-*value calculated with a paired Student’s t-test. *Right:* Marrow to serum ratio was calculated for each individual mouse. Bar represent mean ± SEM. N=6 mice. (B) 1-week-old mouse FFPE femurs were sectioned at 5 µm and subjected to RNA hybridization with a probe specific to *Acp5*, red. Nuclei, blue. GP, growth plate. TB, trabecular bone. BM, bone marrow. White arrowheads indicate multinucleated osteoclasts. Imaged at 20x, shown is a maximum projection of a 3 µm Z-stack, with a step size of 1 µm. (C) Femurs from 5-6 week old C57BL/6J mice were decalcified and paraffin embedded. Sections were hybridized with probes specific to *Acp5* (red) and *Adrb2* (green) to identify osteoclasts expressing *Adrb2*. Nuclei, blue. Shown are representative of results from N=3 separate mice. TB, trabecular bone. BM, bone marrow. White arrowheads indicate multinucleated osteoclasts with green *Adrb2* puncta. Single Z-plane, imaged at 63x.

## Discussion

Here we show that propranolol and intermittent PTH co-treatment improves bone microarchitecture beyond the levels achieved by intermittent PTH treatment alone, which is consistent with previous co-treatment studies ^(15)^. We add to this, however, the novel findings that propranolol enhances tissue mineral density, which may be through promoting Ca^2+^ signaling in response to PTH in osteoblasts. We are also the first to show that propranolol prevents PTH-induced resorption and that this may be through direct inhibition of osteoclast activity, since *Adrb2* is expressed in osteoclasts *in vivo*. This finding in particular suggests more detailed studies examining the roles of βARs in osteoclasts, as well as the impact of βAR-targeting drugs, are critically needed. In the absence of β-blockers, intermittent PTH treatment promotes both bone formation *and* resorption, which could be of concern to individuals with severely lowered bone density and/or high levels of resorption. Intracortical remodeling, which may lead to fracture, has been associated with serum PTH levels in humans, as well as with daily intermittent PTH treatment in rabbits ^(25,26)^. Nonetheless, PTH is very effective at building new bone, so bone loss that occurs from heightened resorption needs to be weighed against its anabolic activity. Our findings suggest that combining PTH with propranolol, or potentially other β-blockers, may be an attractive option to prevent unwanted bone loss in patients with very low BMD.

The mechanism of propranolol action on bone to prevent resorption may be in part through direct effects of propranolol on osteoclasts. Neither PTH-induced serum RANKL nor *Rankl* gene expression was suppressed by propranolol, despite clear improvement in trabecular microarchitecture of the distal femur and the suppressed markers of resorption (serum CTx, *Acp5* expression, and N.Oc/T.Ar, Figure 6). The majority of publications point toward osteoblast expression of RANKL as being the major mediator in the effects of the sympathetic nervous system on osteoclasts ^(2,10)^. However, our findings suggest that this is not the same mechanism through which propranolol blocks resorption. In studies examining the osteoblast-mediated mechanism of SNS-induced bone loss, some experiments suggested no effect of *Adrb2* in osteoclasts. For example, Elefteriou et al. elegantly showed that the non-specific βAR agonist isoproterenol only induces osteoclast differentiation in osteoblast and bone marrow macrophage (BMM) co-cultures when wildtype, but not *Adrb2*^*-/-*^, osteoblasts are present ^(2)^. Similarly, the effect of isoproterenol to induce osteoclast differentiation is not dependent upon the genotype of the BMMs ^(2)^. These data, in combination with findings from models with osteoblast-specific deletions of *Adrb2*, have led to the generally accepted tenet that osteoblasts are the sole mediators of the SNS effect on bone, including the mediators of osteoclast recruitment ^(10)^.

There is a relative scarcity of literature examining the effects of β-adrenergic signaling directly in osteoclasts. However, Rodrigues, et al. showed that propranolol prevented osteoclastogenesis of RAW 264.7 cells through suppression of NFATc1 protein expression, as well as later markers of differentiation: *Acp5, Ctsk*, and *Mmp9* ^(12)^. This suggests a role for βAR signaling in controlling NFATc1, which is under the control of calcium-calmodulin and calcineurin signaling. Kondo, et al. demonstrated that isoproterenol enhanced osteoclastogenesis in bone marrow macrophages and RAW 264.7 cells by increasing reactive oxygen species ^(11)^. Our findings appear more aligned with these reports because we see expression of *Adrb2* in osteoclasts *in vivo* and propranolol-mediated suppression of resorption downstream of osteoblast-induced RANKL. Although these findings are suggestive of a role for β2AR in osteoclasts, additional studies are needed to establish this mechanism *in vivo* and *in vitro*, and to test whether human osteoclasts would have similar properties.

Despite the paucity of literature relating to the effects of the SNS in osteoclasts directly, there is extensive evidence that *Adrb2* is expressed in other cells of the myeloid lineage and that the SNS plays a role in their function ^(27)^. Indeed, the SNS is important for the innate immune response required for tissue repair but can be pathologically activated during certain conditions, such as heart failure. β-blockers are a common therapy for heart failure and work in part by reducing inflammatory cytokine levels ^(27)^. Systemic suppression of inflammation may also be an interesting, yet unexplored, avenue to investigate the mechanism of efficacy of β-blockers in improving BMD, especially with regards to resorption.

In contrast to our findings, global knockout of β2AR prevents any anabolic action of PTH by preventing the PTH-induced increase in mineral apposition and osteoclast number ^(16)^. Although these findings are paradoxical in some sense, they suggest that a basal level of β2-adrenergic signaling in bone may be required for PTH action. It is also clear that pharmaceutical inhibition of β2AR with propranolol is mechanistically distinct from the disruption of signaling through genetic deletion. Propranolol can act as an inverse agonist, meaning it has effects inhibitory toward cAMP signaling even when no agonist is present, reducing spontaneous β2AR receptor activity^(28)^. Furthermore, propranolol also binds to the lesser expressed β1AR and is known to have non-specific effects, including acting as a weak antagonist to some serotonin receptors, which could, in turn, impact bone remodeling^(29,30)^. We cannot exclude the possibility that propranolol is having non-specific effects in our studies, however, we did measure propranolol exposure in serum and marrow and found serum levels comparable to those achieved in patients (Figure 7A), suggesting clinical relevance^(31)^. The published binding affinity of propranolol at the β2AR is 0.5 nM, significantly less than the 107.7 nM achieved in bone marrow (Figure 7A), thus we are clearly inhibiting β2AR with propranolol *in vivo*^(32)^. Interestingly, the binding affinity of NE is 300 nM, while we were measuring levels in the 1-3 nM range in serum (Figure 6E). Collectively, these outcomes suggest that our observations are a result of direct effects of propranolol on β2AR on osteoblasts and osteoclasts, either through competitive inhibition of NE binding, or through inverse agonist properties (of which, IC_50_ values are in the range of the binding affinity)^(28)^.

Moriya et al. showed that *Adrb2* knockdown enhances PTH-induced phosphorylation of CREB, suggesting a suppressive function of β2AR on PTH action ^(21)^. This is consistent with our findings that PTH effects in osteoblasts are enhanced with pharmacological β2AR inhibition. Some intracellular scaffolding proteins, such as members of the Na^+^/H^+^ exchange regulatory cofactor (NHERF) family, selectively promote coupling of the PTH1R to the Gα_q_-PLC-β signaling pathway ^(19)^. NHERF also couples β2AR to the cytoskeleton, and agonist binding causes uncoupling from the cytoskeleton ^(33)^. It is unknown, however, whether propranolol might uncouple β2AR from NHERF, G proteins and/or the other signaling components, which could lead to the enhanced PTH-induced Ca^2+^ flux (Figure 5). Nonetheless, we observed enhanced *Igf1* expression after co-treatment in osteoblasts, which has previously been shown to work in an autocrine manner to stimulate aerobic glycolysis in osteoblasts in response to PTH-treatment at 6 hours ^(24)^. In our hands, the elevation of *Igf1* after 1 hour of propranolol treatment suggests an acceleration of this timetable, which may lead to the improved PTH effect on bone formation and tissue mineral density that we observed *in vivo*. Further studies would be required to confirm this mechanism.

TMD in the primary spongiosa was significantly higher with PTH and propranolol co-treatment compared to PTH alone (Figure 2M). A similar trend was seen with cortical TMD (Figure 3F) as well as total areal BMD assessed by DXA (Table I). Serum P1NP also indicated global increases in bone formation with the combined treatment (Figure 4A), but this was not observed in dynamic histomorphometric measurements of mineral apposition or bone formation in the trabecular bone of the L5 vertebra (Figure 4C). However, gene expression in cortical bone of the tibia suggested more mature osteoblasts/osteocytes (Figure 4G, H), indicating that propranolol may have site-specific effects. Further studies investigating the mineral properties of bone exposed to both PTH and propranolol would be necessary to determine if this combination of treatments may improve strength in patients and animal models.

Clinical literature suggests that β-blockers prevent bone loss and reduce fracture risk in humans. Yang et al. reported that β-blocker use was associated with a 17% decrease in risk of fractures ^(13)^. Toulis et al. analyzed 16 studies involving over 1.5 million subjects and found that the risk of fracture was significantly reduced (by 15%) in subjects receiving β-blockers as compared to control subjects, independent of sex, fracture site, or dose ^(14)^. In contrast to what is the apparent mechanism in rodent models, β1-selective blockers were found to be more effective at reducing fractures than non-selective β-blockers (i.e. propranolol). More recently, when evaluating adrenergic receptor mRNA expression in human bone biopsies, Khosla et al. found that ADRB1 and ADRB2, but not ADRB3, were expressed in human bone ^(34)^. Additionally, patients treated with β1-selective blockers had better bone microarchitecture by quantitative computed tomography (qCT) than non-users, had reduced CTx levels compared to placebo-treated patients, and had increased BMD at the ultradistal radius, but these results were not seen in propranolol (non-selective β-blockers) treated patients. Overall, this data suggests that β1-selective blockers may be more protective against decreased BMD and increased risk of fracture than non-selective β-blockers. Whether the perceived difference between rodents and humans is due to β1AR being understudied in rodent models or due to an actual species-specific difference in the relative importance of β1AR vs β2AR in bone remains unclear.

Our findings reinforce the work of others suggesting that β-blockers, which are routinely used clinically, may be an effective treatment or co-treatment for osteoporosis. Importantly, here we showed that propranolol prevented resorption without modulating PTH-induced RANKL, suggesting that the current understanding of the mechanism of β-blocker and sympathetic nervous system action on bone is incomplete. We have clear evidence of *Adrb2* expression in osteoclasts *in vivo*, and future work should further investigate the extent to which osteoclasts specifically mediate the impact of the SNS on bone. Investigating these mechanisms will be necessary for the accurate interpretation of clinical and preclinical studies examining how the sympathetic nervous system modulates bone homeostasis and contributes to bone pathologies.

## Methods

### Study Approval

The Institutional Animal Care and Use Committee of the Maine Medical Center Research Institute approved all mouse protocols.

### Mice and drug treatment

Sixteen-week-old C57BL/6J (Jackson Laboratories, Bar Harbor, ME) female, intact mice underwent baseline body weight measurements and were randomly assigned to one of four treatment groups. Mice were housed 3-4 per cage and provided standard laboratory chow *ad libitum*. Female mice were chosen because of their higher propensity to have an anabolic response to intermittent PTH. Mice were treated with either vehicle, PTH, propranolol or both PTH and propranolol. PTH (H-1660, Bachem) was aliquoted and stored in glass vials, under argon gas, at −70°C as a 10^−4^ M stock in 4 mM HCl supplemented with 0.1% bovine serum albumin. PTH was thawed and diluted in 0.9% saline immediately prior to injection. Vehicle and PTH (80 μg/kg) were administered by subcutaneous injection five days per week for four weeks. Propranolol hydrochloride oral solution (West-Ward Pharmaceuticals) was dissolved in drinking water at a concentration of 0.5 mg/mL and was delivered daily for four weeks to animals in the propranolol and PTH + propranolol groups as described ^(4)^. At 20 weeks-of-age, mice were sacrificed one hour after treatment and tissues were either fixed in 10% buffered formalin or snap-frozen in liquid nitrogen and stored at −70°C. A similar study was conducted with just 5 days of treatment to obtain additional samples for some mRNA and serum analyses and are identified in the figure legends.

For the detection of propranolol levels in the plasma and bone marrow, sixteen-week-old B6 female mice were administered propranolol dissolved in drinking water as described above. Six mice were separated into three individual cages (two mice/cage). Mice were treated for four days, and sacrificed in the morning, one hour after exchanging treated water for untreated water. Serum was isolated from whole blood and bone marrow was collected from the tibia via centrifugation. Samples were kept frozen until analysis was performed; propranolol levels were detected using LC/MS-MS (see methods below).

### Dual-energy X-ray absorptiometry (DXA)

Mice underwent DXA measurements at baseline and at 20 weeks of age for fat-free mass, fat mass, bone mineral density, and bone mineral content using the PIXImus dual-energy X-ray densitometer (GE-Lunar, Madison, WI). The instrument was calibrated daily with a mouse phantom provided by the manufacturer. Mice were anesthetized with isoflurane and placed ventral side down with each limb and tail positioned away from the body. Full body scans were obtained, and X-ray energy absorptiometry data were gathered and processed with manufacturer-supplied software (Lunar PIXImus 2, version 2.1). The head was specifically excluded from all analyses due to concentrated mineral in skull and teeth.

### Micro-computed tomography (µCT)

A high-resolution desktop micro-tomographic imaging system (µCT40, Scanco Medical AG, Bruttisellen, Switzerland) was used to assess the microarchitecture of the fifth lumbar (L5) vertebral body and the femur in accordance with published guidelines ^(35)^. All analyses were performed using the Scanco Evaluation software.

In the L5 vertebra, scans were acquired using a resolution of 10 μm^3^, 70 kVp peak x-ray tube potential, 114 mA x-ray tube current, 200 ms integration time, and were subjected to Gaussian filtration and segmentation. The endocortical region of the vertebral body was manually selected beginning 100 µm inferior to the cranial growth plate and extending to 100 µm superior to the caudal growth plate. Trabecular bone was segmented from soft-tissue using a threshold of 460 mgHA/cm^3^. Measurements included trabecular bone volume fraction (Tb.BV/TV, %), trabecular thickness (Tb.Th, mm), trabecular number (Tb.N, mm^-1^), trabecular separation (Tb.Sp, mm), connectivity density (Conn.D, mm^-3^), and trabecular bone mineral density (Tb.BMD, mgHA/cm^3^).

To address microarchitecture in the primary and secondary spongiosa of the distal femur and the cortical bone of the femur midshaft, scans were acquired with the same parameters used for the L5 vertebrae. The primary spongiosa region of interest started at the peak of the distal growth plate and extended proximally for 500 µm (50 transverse slices) and included the whole cross-section of the bone. The secondary spongiosa region of interest started immediately superior to the primary spongiosa region and extended 1500 µm (150 transverse slices) proximally and included the endocortical region of the bone. In both regions, bone was segmented from soft-tissue using a mineral density threshold of 460 mgHA/cm^3^. Trabecular bone in the endocortical area of the secondary spongiosa region was analyzed for bone volume fraction (Tb.BV/TV, %), trabecular thickness (Tb.Th, mm), trabecular number (Tb.N, mm^-1^), trabecular separation (Tb.Sp, mm), connectivity density (Conn.D, mm^-3^), and trabecular bone mineral density (Tb.BMD, mgHA/cm^3^). The bone in the primary spongiosa was analyzed for bone volume, total volume, and bone mineral density.

Cortical bone architecture was analyzed in a 500 µm long region at the femoral mid-diaphysis (55% of the total femur length inferior to the top of the femoral head). Cortical bone was segmented from soft tissue using a threshold of 700 mgHA/cm^3^. Measurements included cortical bone area (Ct.Ar, mm^2^), total cortical cross-sectional area (Tt.Ar, mm^2^), cortical bone area fraction (Ct.Ar/Tt.Ar, %), cortical thickness (Ct.Th, mm) and cortical porosity (%).

### Histology and Histomorphometry

Tibiae were fixed in 10% neutral-buffered formalin and transferred to 70% ethanol after 24 hours. Samples were decalcified, paraffin-embedded, sectioned, and stained with hematoxylin and eosin. Adipocyte size and number in the marrow in the region of the secondary spongiosa were analyzed with BIOQUANT OSTEO software (BIOQUANT Image Analysis Corporation, Nashville, TN). Total adipocyte area was divided by total area (marrow plus trabecular bone) in the secondary spongiosa and multiplied by 100 to calculate adipocyte volume / total volume (AV/TV). Adipocyte number was normalized to the total area measured.

Vertebrae were fixed in 10% neutral-buffered formalin and transferred to 70% ethanol after 24 hours. The fixed lumbar vertebrae (L2-L5) were dehydrated with acetone and embedded in methylmethacrylate. Undecalcified 4-µm-thick sections were obtained by microtome and stained with Von Kossa method for showing the mineralized bone. The consecutive second section was left unstained for the analysis of fluorescence labeling and the third section was stained with 2% Toluidine Blue (pH 3.7) for the analysis of osteoblasts, osteoid, and osteoclasts. The bone histomorphometric analysis was performed in the lumbar vertebra under 200X magnification in a 1.35 mm high x 1.3 mm wide region where was located 400 μm away from the upper and lower growth plate using OsteoMeasure analyzing software (Osteometrics Inc., Decatur, GA), in accordance with published guidelines ^(36)^. The structural parameters (BV/TV, Tb.Th, Tb.N, and Tb.Sp) were obtained by taking an average from two different analysis of consecutive sections. The structural, dynamic and cellular parameters were calculated and expressed according to the standardized nomenclature ^(36)^.

### Enzyme Immunoassays

Serum concentrations of amino-terminal propeptide of type 1 procollagen (P1NP), cross-linked C-telopeptide (CTx) and TNF-related activation-induced cytokine (RANKL) were measured with the Rat/Mouse P1NP enzyme immunoassay (EIA), RatLaps EIA (both from Immunodiagnostic Systems, Scottsdale, AZ), and the Quantikine ELISA Mouse TRANCE/RANKL/TNSFSF11 Kit (R&D Systems, Minneapolis, MN). The intraassay variations were 6.3%, 6.9% and 4.3%, and the interassay variations were 8.5%, 12% and 6.9% respectively. All measurements were performed in duplicate.

### Osteoblast assays

MC3T3-E1 (clone 4) cells were purchased from ATCC and maintained at a low passage number in α-MEM containing 1% penicillin streptomycin (P/S) and 10% fetal bovine serum (FBS). For calcium assays, cells were plated overnight in 96-well black wall plates at a density of 5×10^4^ cells/well. Cells were then differentiated for 7 days with ascorbic acid and β-glycerophosphate as previously described ^(37)^. PTH was aliquoted and stored as above, and thawed and diluted immediately prior to use. Propranolol hydrochloride (P0884, Sigma-Aldrich) was dissolved in 1x HBSS buffer containing 20 mM HEPES for each experiment. Calcium assays were performed with the FLIPR Calcium 6 Assay Kit (Molecular Devices) according to the manufacturer’s protocol. In brief, on day 7 of differentiation, cells were incubated with calcium dye for two hours at 37°C and 5% CO_2_. Then, fluorescence was measured on a FlexStation 3 Multi-Mode Microplate Reader using SoftMax Pro 7 software (Molecular Devices) in kinetic mode in response to additions of a range of concentrations of PTH. Change in relative fluorescence units (maximum – minimum) is indicative of intracellular calcium (iCa^2+^) levels. Concentration-dependent change in max-min value was plotted on a logarithmic scale and fit with a 4-parameter logistic curve to calculate the EC_50_. In some experiments, calcium dye incubation was concurrent with pretreatment with propranolol. Additional 24-well plates were treated simultaneously with PTH and propranolol for 1 hour, without the Calcium 6 reagents, and were saved for mRNA analyses.

### Propranolol extraction and measurements

Marrow was prepared for extraction by homogenization at a 1:1 ratio with 50 mM potassium phosphate buffer pH 7.4. A volume of 20 µL of serum or marrow homogenate was combined with 100 µL of HPLC grade acetonitrile and vortex mixed for two minutes. Subsequent to centrifugation at 14000 rpm for five minutes, the supernatant was transferred to a 96-well plate for liquid chromatograph tandem mass spectrometry (LC/MS-MS) analysis. A calibration curve was formed in mouse plasma from 1.00-200 nM by serial dilution and extracted via the same methodology. An Agilent 1200 system consisting of a binary pump, column compartment and autosampler was used for solvent delivery and sample introduction. Chromatographic separation was performed on an Agilent Zorbax SB C18 2.1 x 50 mm 2.7 µm column via a gradient using 0.1% formic acid in water (A) and 0.1% formic acid in acetonitrile (B). Gradient elution was 95% A ramping to 5% A from 0.0-3.0 minutes, with re-equilibration at initial conditions from 3.0 to 4.0 minutes. The flow rate was 0.75 mL/min, and column temperature was 30°C. Detection of propranolol was obtained using an Agilent 6460 triple quadrupole mass spectrometer, monitoring the transition 260.0 → 116.0 with a fragmentor of 86 V and a collision energy of 21 V. The retention time of propranolol was 1.89 minutes.

### RNA isolation and real-time PCR (qPCR)

Total RNA was isolated from whole tibia and cortical shell using the standard TRIZOL (Sigma, St. Louis, MO) method. Total RNA from cells was isolated using an RNeasy Mini Kit (Qiagen). cDNA was synthesized using the High Capacity cDNA Reverse Transcriptase Kit (Applied Biosystems, Foster City, CA) according to the manufacturer’s instructions. mRNA expression analysis was performed using an iQ SYBR Green Supermix or Taqman Gene Expression Assays with an iQ5 thermal cycler and detection system (Bio-Rad, Hercules, CA). Hypoxanthine guanine phosphoribosyl transferase (*Hprt*) was used as an internal standard control gene^(38)^. Primer sequences are listed in the supplemental information (Table S1).

### Norepinephrine extraction and measurements

A volume of 20 µL of serum was combined with 100 µL of HPLC grade acetonitrile and vortex mixed for two minutes. Subsequent to centrifugation at 14000 rpm for five minutes, the supernatant was transferred to a 96-well plate for liquid chromatograph tandem mass spectrometry (LC/MS-MS) analysis. A calibration curve was formed in mouse plasma from 0.305-2500 nM by serial dilution and extracted via the same methodology. An Agilent 1200 system consisting of a binary pump, column compartment and autosampler was used for solvent delivery and sample introduction. Chromatographic separation was performed on a Phenomenex Hydro RP 2.0 x 150 mm 4 µm column via a gradient using 0.1% formic acid in water (A) and 0.1% formic acid in acetonitrile (B). Gradient elution was 98% A from 0-1 minute, ramping to 50% A from 1.1 to 3.0 minutes, holding at 50% until 5.5 minutes, with re-equilibration at initial conditions from 5.6 to 7.5 minutes. The flow rate was 0.30 mL/min, and column temperature was 30°C. Detection of norepinephrine was obtained using an Agilent 6460 triple quadrupole mass spectrometer, monitoring the transition 152.0 → 107.0 with a fragmentor of 94 V and a collision energy of 18 V. The retention time of norepinephrine was 1.37 minutes.

### Detection of Adrb2 mRNA transcript in femur

Femurs were isolated from 1-week-old or 6-week-old B6 mice and fixed in 10% neutral buffered formalin for 24 hours, then decalcified in 0.5M EDTA pH 7.8 solution by rocking at 4LJ C for 1-2 weeks. Samples were dehydrated in 70% EtOH, embedded in paraffin, and sectioned in 5 μM sections using standard methods. RNA *in situ* hybridization was performed using the RNAscope FFPE Multiplex Fluorescent Assay V2 (ACD Bio) per manufacturer’s instructions. Briefly, sections were deparaffinized followed by pre-treatment using hydrogen peroxide and target retrieval was achieved by steaming for 15 minutes. Tissue sections were treated with protease, and the RNAscope assay was performed as outlined by manufacturer’s instructions using probes for *Acp5* and *Adrb2*. Images were acquired using a Leica TCS SP8 laser scanning confocal microscope with a 20x air objective or 63x oil objective.

### Sample size estimation and statistics

Sample size for mouse experiments was estimated based on previous experiments in our laboratory using trabecular BV/TV as a primary outcome. Secondary outcomes of interest included serum and histomorphometric indices of bone remodeling. All statistics were performed with Prism 7 statistical software (GraphPad Software, Inc., La Jolla, CA). Normally distributed data were analyzed for statistically significant differences using either a two-sided Student’s *t*-test or a two-way ANOVA followed by Holm-Sidak multiple comparison *post hoc* test, where appropriate. Statistical significance was set at *p* < 0.05. All data are expressed as the mean ± standard error of the mean (SEM). Outliers were handled as follows: first, any scientific/biological reason for the outlier was investigated (i.e. sick mouse, degraded RNA, bad histology section); second, if no explanation for the outlier was found we performed an outlier test to determine if the datum was > 3 standard deviations (SD) from the mean of the other samples in that group. If the datum was > 3 SD from the mean, it was excluded from further analysis. Outlier handling did not significantly change the interpretation of the results.

## Author Contributions

AT was responsible for experimental design, data acquisition, data analysis, interpretation, and drafting of the manuscript. ACB was responsible for experimental design, data acquisition, data analysis, interpretation, drafting and critical review of the manuscript. DJB was responsible for data acquisition, data analysis, interpretation, drafting of and critical revision of the manuscript. ALL was responsible for experimental design, data acquisition, data analysis, interpretation and drafting and critical review of the manuscript. HH was responsible for data acquisition, data analysis, interpretation and drafting of the manuscript. KTN was responsible for data analysis, drafting and critical review of the manuscript. KN was responsible for data acquisition, data analysis, interpretation, drafting of and critical revision of the manuscript. DB was responsible for experimental design, data acquisition, interpretation, drafting and critical revision of the manuscript. KLH was responsible for experimental design, data interpretation, drafting and critical revision of the manuscript. RB was responsible for data interpretation and critical revision of the manuscript. MLB was responsible for data interpretation and critical revision of the manuscript. ARG was responsible for experimental design, data interpretation and critical revision of the manuscript. KJM was responsible for experimental design, data acquisition, data analysis, interpretation, drafting and critical revision of the manuscript. All authors had final approval of the manuscript.

## Acknowledgements

The authors thank Adriana Lelis Carvalho and Ryan Neilson for critical evaluation of the work. This work was supported by the National Institute of Arthritis And Musculoskeletal And Skin Diseases (NIAMS) and the National Institute of General Medical Sciences (NIGMS) of the National Institutes of Health (NIH) under award numbers K01AR067858 and P20GM121301 to KJM. This work utilized services of the Maine Medical Center Research Institute (MMCRI) Molecular Phenotyping Core, which is supported by NIH/NIGMS P30GM106391, the Physiology Core, which is supported by NIH/NIGMS P30GM106391 and P20GM121301, the Histopathology and Histomorphometry Core, which is supported by NIH/NIGMS P30GM106391, P20GM121301, and P30GM103392, and the Mouse Transgenic and In Vivo Imaging Core which is supported by NIH/NIGMS P30GM103392. All cores also received support from the Norther New England Clinical and Translational Research Network NIH/NIGMS U54GM115516. The content is solely the responsibility of the authors and does not necessarily represent the official views of the National Institutes of Health.

---

**Figure S1.**
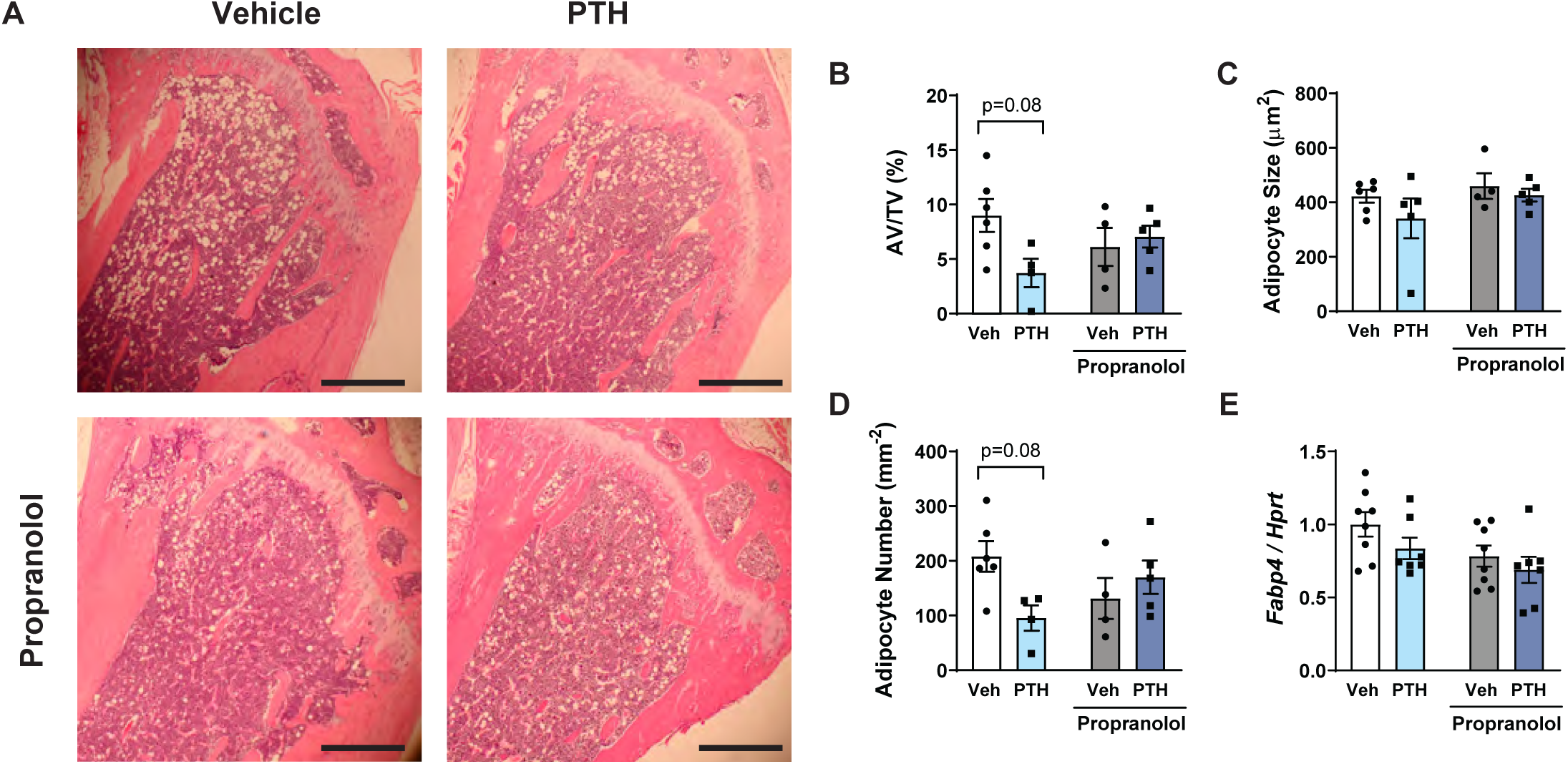
Effects of PTH and propranolol on proximal tibia bone marrow adiposity. Mice were treated for 4 weeks with vehicle (white), 80 µg/kg PTH (light blue), 0.5 mg/ml propranolol (gray), or PTH and propranolol (dark blue). (A) Tibias were decalcified and stained with H & E. Scale bar = 500 µm. (B-D) Adipocyte ghosts were measured using BIOQUANT OSTEO software. (E) *Fabp4* gene expression was measured in whole tibia and normalized to non-modulated housekeeping gene *Hprt*. Bars represent mean ± standard error. Indicated *p*-values were determined by Holm-Sidak *post hoc* test after a significant interaction effect was found using two-way ANOVA.

**Figure S2.**
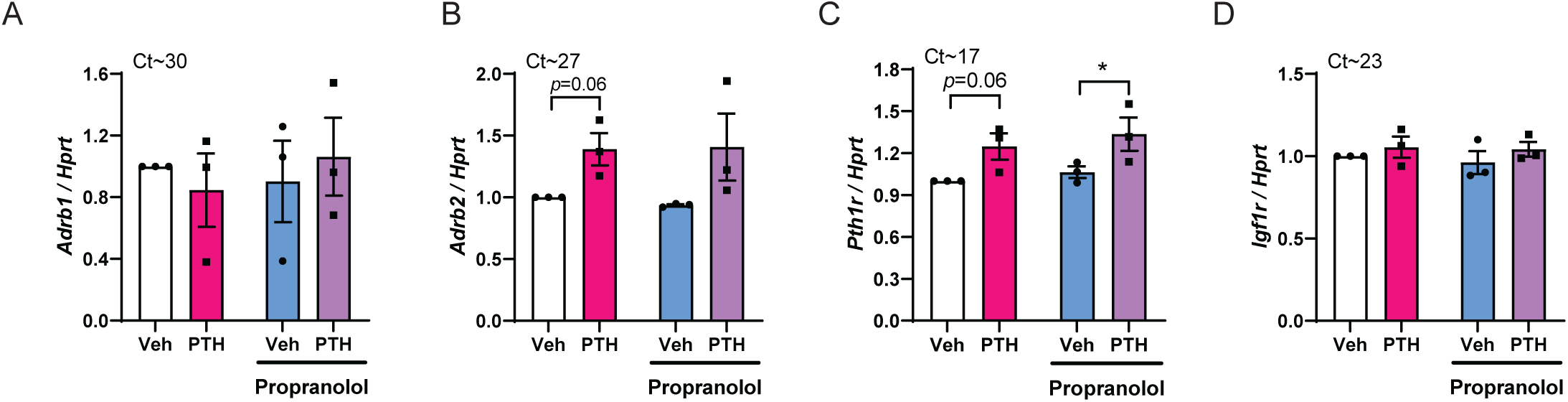
Relevant receptor expression levels in osteoblast-like cells. MC3T3-E1 cells were differentiated to day 7 in osteogenic media. Gene expression of *Adrb1, Adrb2, Pth1r* and *Igf1r* in MC3T3-E1 cells at day 7 after 1 hour treatment with vehicle (white), 100 nM PTH (pink), 1 µM propranolol (blue) or 100 nM PTH + 1 µM propranolol (purple). Genes are normalized to non-modulated housekeeping gene, *Hprt*. Each data point represents the mean from an independent experiment, performed in triplicate. Expression levels in vehicle-treated cells from each experiment were set to 1. Approximate Ct values of vehicle-treated groups are shown above each graph. Bars represent mean of the three independent experiments ± SEM. \**p*<0.05, \*\**p*<0.01 by Holm-Sidak *post hoc* test after a significant two-way ANOVA.

