## Supplemental Table I for "Propranolol promotes bone formation and limits resorption through novel mechanisms during anabolic parathyroid hormone treatment in female C57BL/6J mice"

Table SI. qPCR primer information.

| **Target Gene** | **Source/Supplier** | **Sequence** | **Catalog Number** | **Reference** |
| --- | --- | --- | --- | --- |
| ***Col1a1*** | Primer Design (Southampton, UK) | Forward: 5'-TCG TGG CTT CTC TGG TCT C-3' | N/A | N/A |
|  |  | Reverse: 5'-CCG TTG AGT CCG TCT TTG C-3' |  |  |
| ***Ctsk*** | IDT (Coralvile, IA) | Forward: 5’-GCA GAG GTG TGT ACT ATG-3’ | N/A | N/A |
|  |  | Reverse: 5’-GCA GGC GTT GTT CTT ATT-3’ |  |  |
| ***Dmp1*** | IDT (Coralvile, IA) | Forward: 5'-TCG CTG AGG TTT TGA CCT TGT-3' | N/A | ^(1)^ |
|  |  | Reverse: 5'-CTC ACT GTT CGT GGG TGG TG-3' |  |  |
| ***Fabp4*** | IDT (Coralvile, IA) | Forward: 5′-GCG TGG AAT TCG ATG AAA TCA-3′ | N/A | ^(2)^ |
|  |  | Reverse: 5′-CCC GCC ATC TAG GGT TAT GA-3′ |  |  |
| ***Fos*** | Qiagen (Germantown, MD) | Not provided | 330001, PPM02940C | N/A |
| ***Hprt*** | IDT (Coralvile, IA) | Forward: 5′-AAG CCT AAG ATG AGC GCA AG-3' | N/A | ^(3)^ |
|  |  | Reverse: 5′-TTA CTA GGC AGA TGG CCA CA-3′ |  |  |
| ***Igf1*** | Primer Design (Southampton, UK) | Forward: 5’-GAC CGA GGG GCT TTT ACT TC-3’ | N/A | N/A |
|  |  | Reverse: 5'TGC TTT TGT AGG CTT CAG TGG-3’ |  |  |
| ***Opg*** | IDT (Coralvile, IA) | Forward: 5’GAAGAAGATCATCCAAGACATTGAC-3’ | N/A | ^(4)^ |
|  |  | Reverse: 5’-TCCATAAACTGAGTAGCTTCAGGAG-3’ |  |  |
| ***Osteocalcin (Bglap)*** | IDT (Coralvile, IA) | Forward: 5'-ACG GTA TCA CTA TTT AGG ACC TGT-3' | N/A | ^(5)^ |
|  |  | Reverse: 5'-ACT TTA TTT TGG AGC TGC TGT GAC-3' |  |  |
| ***Pth1r*** | IDT (Coralvile, IA) | Forward: 5'-TTT CCC GGT GCC TTC TCT TTC-3’ | N/A | ^(6)^ |
|  |  | Reverse: 5’-CAG GCG CAA TGT GAC AAG C-3’ |  |  |
| ***Rankl*** | Primer Design (Southampton, UK) | Forward: 5'-TTT GCA CAC CTC ACC ATC AAT-3' | N/A | N/A |
|  |  | Reverse: 5'- CCC TTA GTT TTC CGT TGC TTA AC-3' |  |  |
| ***Runx2*** | IDT (Coralvile, IA) | Forward: 5'-GAC AGA AGC TTG ATG ACT CTA AAC C-3' | N/A | ^(7)^ |
|  |  | Reverse: 5'-TCT GTA ATC TGA CTC TGT CCT TGT G-3' |  |  |
| ***Sost*** | Qiagen (Germantown, MD) | Not provided | 330001, PPM36047A | N/A |
| ***Trap5 (Acp5)*** | IDT (Coralvile, IA) | Forward: 5'-AAT GCC TCG ACC TGG GA-3' | N/A | ^(8)^ |
|  |  | Reverse: 5'-CGT AGT CCT CCT TGG CTG CT-3' |  |  |
